# Chronic obesity does not alter cancer incidence in *Trp53*^*R270H/*+^ mice

**DOI:** 10.1101/2024.10.14.618190

**Authors:** Ilaria Panzeri, Zachary Madaj, Luca Fagnocchi, Stefanos Apostle, Megan Tompkins, Galen Hostetter, J. Andrew Pospisilik

## Abstract

Obesity is a complex chronic disease characterized by excessive adiposity and associations with numerous co-morbidities, including cancer. Despite extensive research, we have limited understanding of the mechanisms coupling obesity to cancer risk, and separately, of the contexts where obesity does or does not exacerbate disease. Here, we show that chronic high-fat diet induced obesity has no significant effect on the *Trp53*^*R270H/+*^ mouse, a model of human Li-Fraumeni multi-cancer syndrome. Surprisingly, despite inducing rapid and highly penetrant obesity and long-term differences in adiposity, greater than one year of HFD had no significant effect on survival or tumor burden. These findings were replicated in two separate cohorts totaling 359 mice and thus provide important negative data for the field. Given strong publication bias against negative data in the literature, this large murine cohort study represents a clear case where chronic diet-induced obesity does not accelerate or aggravate cancer outcomes. The data thus carry high impact for researchers, funders, and policymakers alike.

## INTRODUCTION

Obesity is a chronic metabolic condition characterized by elevated body weight, adiposity, and numerous comorbidities, including cardiovascular disease, diabetes and autoimmunity. Obesity (and more moderate ‘overweight’) affect ∼1.3 billion individuals globally, and the direct economic costs in the US alone are estimated at $173 billion per year^1^. In addition to cardiometabolic complications, obesity associates with increased incidence and severity of multiple cancers, including at least 13 distinct anatomical sites^2-5^. Obesity has also been associated with altered treatment efficacy and toxicity^6,7^. Further, obesity correlates with higher recurrence rates and poorer cancer prognoses^8^, findings that have led to the inclusion of weight management in clinical guidelines for cancer survivors^9-11^.

While numerous mechanisms have been proposed to explain how obesity drives cancer^12-16^, deep understanding remains elusive. We currently do not know *how* obesity influences tumorigenesis, *which* cancer types are susceptible to obesity-dependent regulation, or *when* during tumorigenesis such regulation most relevant. This lack of understanding derives from methodological issues, as well as questions of reverse causality, detection and selection biases, and the overall breadth of cancers observed in the clinic^17,18^.

Given these substantial limitations in the state of the art, accurate literature is of utmost importance. Negative findings in biomedical research, despite being a vital part of the research record, are often underreported^19-23^. This specific ‘publication bias’ can carry real-life consequences for patients^24,25,26^ and publication of negative data is therefore essential. This bias is becoming increasingly problematic in the era of artificial intelligence and machine learning, where models trained exclusively on published “positive” results inherit and potentially amplify existing biases, leading to skewed predictions and unrealistic assessments of experimental outcomes^27^. Though equally challenging to address, the origins of publication bias against negative data stem from multiple factors: (i) insufficient professional incentives to publish negative findings, (ii) a research culture that undervalues negative results, and (iii) perceived low interest from editors and reviewers (though this last factor has proven less widespread than commonly assumed)^28-31^. These and other factors, coupled with funding struggles mean that investigators have little to no motivation to complete fully powered negative studies once pilot data suggest weak or absent phenotypes. Indeed, when they do appear, negative data are often buried in the supplements of a manuscript focusing on a positive finding and are thus more likely to be overlooked. With respect to obesity and cancer, recent literature does contain select reports that question the causal association between body mass index (BMI) and reduced cancer survival^32-42^.

This work presented here tested the interaction between chronic high-fat feeding and a *Trp53*-model of spontaneous murine multi-cancer syndrome (MCS) (the *Trp53*^*R270H/+*^ mouse). *TP53* is one of the most commonly mutated tumor suppressor genes in human cancers^43^ and loss-of-function or dominant negative mutations cause Li-Fraumeni syndrome (LFS)^44-46^. LFS is an autosomal dominant disorder characterized by a marked susceptibility to diverse p53-dependent cancers. *Trp53*^*R270H/+*^ mice (the equivalent of the human Li-Fraumeni hotspot mutation R273H) faithfully recapitulate many aspects of the human MCS disorder, exhibiting a broad spectrum of tumors, including a variety of carcinomas, soft tissue and bone sarcomas, leukemia, and even glioblastoma (the most common brain cancer in LFS patients). *Trp53*^*R270H/+*^ mice exhibit a mean lifespan survival of approximately 1 year^46^.

High-fat diet (HFD) feeding is the most commonly used murine model of obesity^47,48^. The utility of the model is underpinned by human epidemiological studies that associate increased dietary fat intake with obesity incidence^49-51^. Elevated dietary fat intake in humans and mice increases adiposity and associated hyperglycemia, inflammation, hypertension, plasma lipids, insulin resistance, and reduced beta-cell function^52^. Over time, this collectively leads to ‘metabolic syndrome’ characterized by elevated adiposity, cardiovascular disease, hepatosteatosis, type 2 diabetes, and increased mortality^53,54^. Importantly, HFD treatment is just one of many murine models of human obesity (and modern Western diet composition). Just as human obesity is increasingly acknowledged as a heterogeneous set of disorders^55,56^ with numerous distinct genetic, epigenetic, and environmental drivers, HFD treatment models only a subset of the many recognized drivers of obesity. HFD has been shown to exacerbate a number of cancer models in mice, including colorectal cancer^57^, hepatocellular carcinoma^58^, pancreatic ductal adenocarcinoma^59^, breast cancer^60^, prostate cancer^61^, and skin carcinogenesis models^62^.

Here, we report a deep longitudinal analysis of two large cohorts of *Trp53*^*R270H/+*^ animals challenged life-long with HFD or control chow diet (CD) from an early age. Surprisingly, despite rapid and highly penetrant induction of obesity and associated metabolic sequelae, we find no effect of HFD on overall survival, tumor burden, or tumor spectrum. The data highlight an important example where chronic obesity does not accelerate or aggravate cancer etiology.

## RESULTS

### A cohort to study chronic effects of obesity on *Trp53*-dependent cancers

We set up two cohorts totaling 359 animals, with equal representation of females and males. The cohorts included 203 *Trp53*^*R270H/+*^ animals and 156 wildtype (WT) littermates to control for potential developmental effects^63^. All mice were fed CD until 8 weeks of age, at which point they were randomly assigned to CD or HFD groups for the remainder of the experiment. We tracked animals from birth to the predetermined endpoint of 70 weeks of age (more than 1 year of HFD) regularly monitoring morphological, growth and metabolic characteristics, as well as performing health checks for signs of cancer onset 2-3 times per week (**Fig.1A**). At sacrifice, all animals underwent a 21-organ dissection protocol in which tissues were isolated, processed for histology, and scored by a board-certified pathologist.

**Figure 1.**
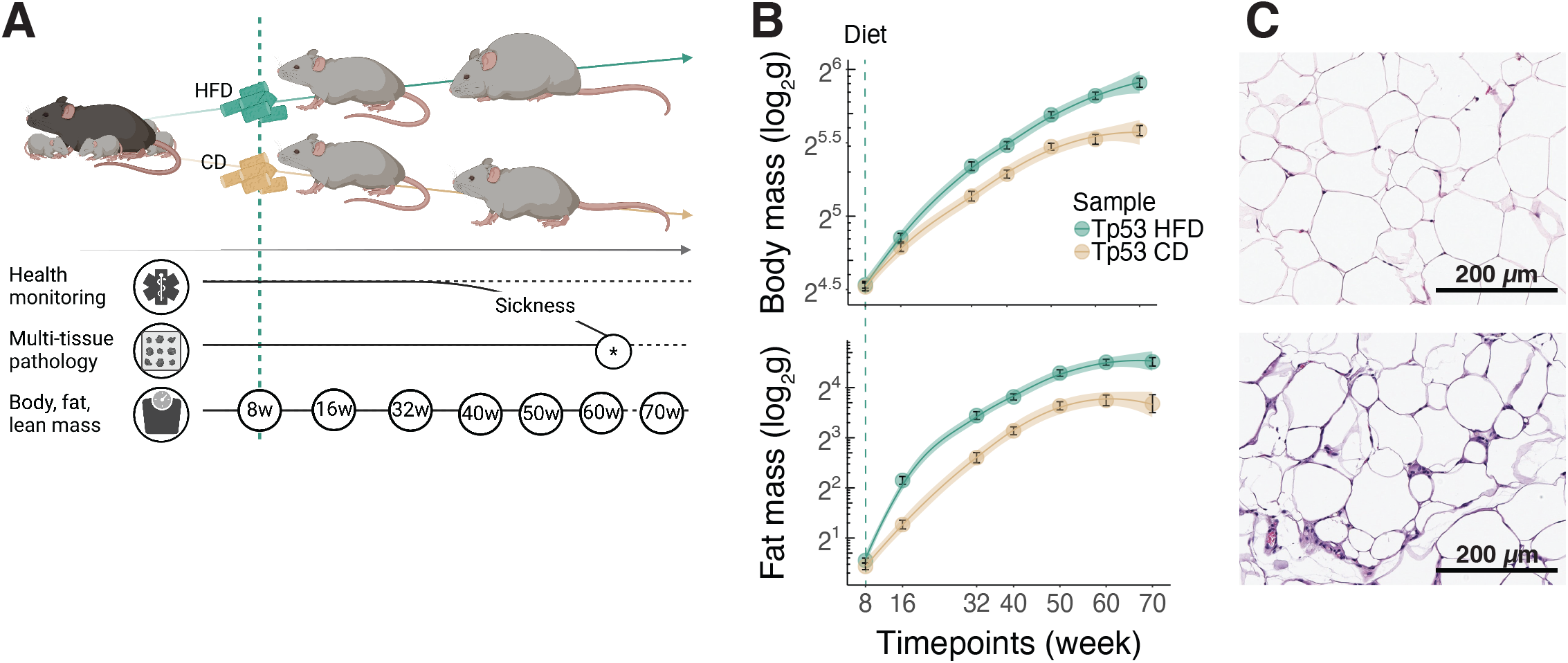
A cohort to study chronic effects of obesity on *Tp53*-dependent cancers. **A)** Schematic of the experimental plan. *Tp53*^*+/R270H*^ females were mated with the *Trim28*^*+/D9*^ *FVB*.*J* males. F1 genotypes were screened for health issues and mass development. Tissues were harvested at sickness report. Histopathology determined the presence of tumors. Body, fat, and lean mass were measured at multiple timepoints. Created with BioRender.com. **B)** Scatter plots and smoothed conditional means (95% confidence interval, “loess” method) for *body* (<u>top</u>) and *fat* (<u>bottom</u>) mass in *Tp53*^*+/R270H*^ females and males (pooled data). N=195 animals (101 females and 94 males). **C)** Representative examples of hematoxylin and eosin-stained adipose tissue from *chow-*(<u>top</u>) and *high-fat* (<u>bottom</u>) diet-fed *Tp53*^*+/R270H*^ male animal at 70 weeks of age. N=1 animal.

Mice of both genotypes and sexes responded rapidly and consistently to the HFD intervention with significant gains in overall fat and whole-body mass, compared to controls (**Fig.1B, Fig.S1A**). Notable given previous reports of metabolic roles for p53^64^, we found no significant differences in HFD-triggered body or fat mass accumulation in *Trp53*^*R270H/+*^ compared to WT animals (**Fig.S1B**). Histopathological evaluation revealed increased immune infiltration, lipoblast activity (indicative of adipose tissue proliferation), adipocyte size and adipocyte shape heterogeneity in HFD-treated animals (**Fig.1C**). Thus, we generated a large-scale cohort for assessing interaction between HFD-induced obesity and *Trp53*^*R270H/+*^ induced cancer. The data demonstrate that the *Trp53*^*R270H/+*^ mutation does not impact the physiological response to HFD.

### HFD does not alter survival in *Trp53*^*R270H/+*^ mice

To our surprise, despite ∼1 year of obesogenic diet and associated obesity, we found no change in survival probability of the animals. *Trp53*^*R270H/+*^ animals under CD or HFD showed near-identical median survival times around 64-65 weeks of age (**Fig.2A**). No evidence of confounding heterogeneity effects was observed (e.g., altered shape of the Kaplan-Meier curve). The negative finding was true for both females and males (**Fig.S2A**), and importantly, held true for two separate discovery and validation cohorts run ∼2 years apart (**Fig.S2B**). One discovery cohort was run and analyzed before the COVID19 pandemic, and the validation cohort after. Both cohorts included at least 14 animals per genotype, sex and diet. Animals from the two cohorts were derived from distinct sets of breeding pairs (arguing against unexpected parental or litter-specific anomalies) and both were challenged with distinct manufacturing batches of HFD/CD. These data demonstrate a reproducible lack of effect of chronic HFD on *Trp53*^*R270H/+*^ survival. Thus, *Trp53*^*R270H/+*^*-*triggered cancer outcome is largely refractory to HFD induced obesity.

**Figure 2.**
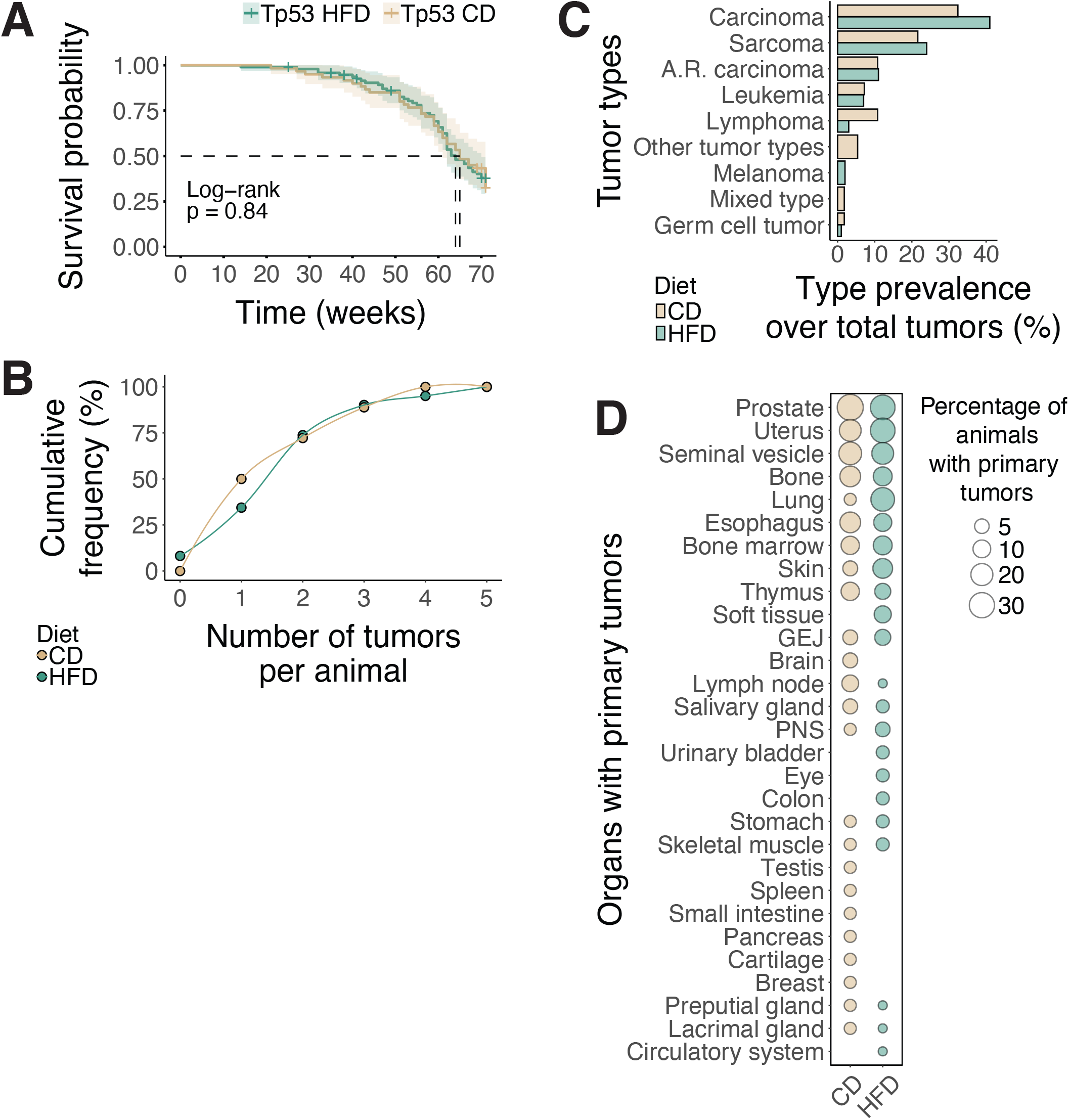
HFD does not alter survival, burden, or spectrum in *Tp53*^*R270H/+*^ mice. **A)** Kaplan-Meier survival probability by diet for *Tp53*^*+/R270H*^ animals. Log-rank test, p=0.84. N=155 animals (95 high-fat diet vs 60 chow diet, pooled female and male data). **B)** Cumulative distribution of tumor burden (number of tumors per animal) in *Tp53*^*+/R270H*^ animals fed with chow– or high-fat diet. No significant differences were observed according to an asymptotic two-sided, two-sample Kolmogorov-Smirnov test (D = 0.020408, p-value = 1). N=97 animals (61 high-fat diet vs 36 chow diet, pooled female and male data). **C)** Prevalence of each tumor type over the total number of tumors in chow– or high-fat diet-fed *Tp53*^*+/R270H*^ animals. N=168 tumors (108 tumors in high-fat diet– vs 60 in chow diet-fed animals, pooled female and male data). **D)** Percentage of *Tp53*^*+/R270H*^ animals with primary tumors targeting the different organs. N=92 animals (61 high-fat diet vs 36 chow diet, pooled female and male data).

### HFD shows minimal impact on tumor burden, prevalence and spectrum in *Trp53*^*R270H/+*^ mice

True to the model, most *Trp53*^*R270H/+*^ animals exhibited high tumor burden with multiple primary tumors per animal. Once again however, HFD animals showed no difference in tumor burden relative to CD animals (**Fig.2B**). This observation held for independent analyses of female and male animals (**Fig.S2C**) and across discovery and validation cohorts (**Fig.S2D**). Thus, chronic HFD-induced obesity does not alter tumor burden in *Trp53*^*R270H/+*^ mice.

As introduced earlier, Li-Fraumeni syndrome and the *Trp53*^*R270H/+*^ model are characterized by heightened cancer incidence across a range of tumor types, and a wide range of targeted tissues. Where sufficiently powered, we also tested for HFD-induced changes in tumor type. Consistent with literature^46^, the majority of tumors were carcinomas and sarcomas, followed by leukemias and lymphomas (**Fig.2C**). Primary sites for tumor emergence included the reproductive system (prostate, uterus, seminal vesicles), bone and bone marrow, lungs, esophagus, skin, thymus, and soft tissues (**Fig.2D**). No evidence of changed tumor type (carcinoma and sarcoma) was found comparing HFD and CD *Trp53*^*R270H/+*^ animals (**Fig.2C**). Again, these observations were true for both female and male comparisons (**Fig.S2E**). The large cohort size notwithstanding, the study was not powered to detect differences in targeted tissues (**Fig.2C, Fig.S2F**).

Thus, *Trp53*^*R270H/+*^-dependent cancers are largely refractory to the effects of HFD-induced obesity.

## DISCUSSION

Overall, our data provide a clear example where obesity and HFD-induced metabolic changes have limited impact on tumorigenesis and cancer outcomes. This finding is significant in the context of known publication bias against negative data^24^. Our study provides empirical support for several epidemiological reports showing little to no association between obesity and select subsets of human cancer types^8,18,65-67^.

Despite extensive evidence linking obesity to heightened cancer risk through mechanisms such as chronic inflammation, insulin resistance, and altered hormonal signaling^52,68^, our *Trp53*^*R270H/+*^ mouse model did not exhibit worsened tumorigenesis under HFD-induced obesity. One possible explanation is that obesity-associated pro-inflammatory effect (especially sterile inflammation) might depend on additional physiological stressors (e.g., oxidative stress, tissue hypoxia, or subclinical infections) that were absent in our controlled conditions^69^. Supporting this hypothesis, certain dietary fats have been shown to induce complement activation and intestinal tumorigenesis independently of obesity, emphasizing dietary composition and inflammation rather than obesity itself as the critical factor^70^. These observations align with our previous findings, where *Trim28*^*+/D9*^ mice exhibited increased adiposity without accompanying inflammatory responses or heightened cancer incidence, suggesting that metabolic health rather than adiposity per se may determine cancer outcomes^55,63^. Moreover, the genetic context of the *Trp53*^*R270H/+*^ mutation itself may alter the tissue microenvironment, potentially buffering inflammatory disruptions typically driven by obesity. Multiple studies identify p53 as a key regulator of adipose tissue inflammation, immune cell infiltration, and adipokine signaling, influencing adipocyte–macrophage crosstalk, insulin sensitivity, and sterile inflammation^71,72^. Indeed, adipose-tissue specific deletion of *Trp53* (*Trp53*^*fl/fl*^; *Fabp4-Cre*) has been shown to normalize insulin glucose tolerance, and cytokine misexpression in obesity-prone agouti mice (Ay) as well as correlate with decreased senescence-like features^73^.

p53 further directly regulates energy metabolism, glycolysis, oxidative phosphorylation, and fatty acid oxidation, which could counterbalance the adverse metabolic and inflammatory profiles associated with high-fat feeding^64,74,75^. Consistent with these observations and in line with related metabolic changes reported in Li-Fraumeni syndrome^76^, partial inhibition of fatty acid oxidation has previously delayed tumor onset in *Tp53*^*R270H/+*^ mice^77-79^.

Collectively, these findings suggest that the metabolic and immune landscape induced by the *Trp53*^*R270H*^ mutation could counteract or buffer the inflammatory and metabolic disruptions typically associated with chronic obesity.

Importantly, while our data clearly show a lack of effect of HFD and HFD-induced obesity on p53-dependent tumorigenesis, they do not rule out all potential associations between obesities *and* R270H-driven cancer. Our limited understanding of the obesity-cancer associations is exacerbated by an underappreciation for the heterogeneity of human obesity itself. Our work^54-56,63^ and that of others^80-82^ have highlighted that obesity is an umbrella term for what are a collection of disorders.

Heterogeneous clinical presentations include metabolically ‘healthy’ and ‘unhealthy’ obesities, Types-A and –B obesities^56^, and a complex intersection of clinically relevant heterogeneity in body size, shape and composition^83-85^. While our work highlights a negative interaction between HFD-induced obesity and *Trp53*^*R270H/+*^ cancers, this does not mean that other obesity ‘endotypes’ (i.e., etiologically distinct obesity subtypes) will not exhibit a strong interaction. Testing causal links between multiple obesity subtypes and their respective dietary associations, across myriad cancer types, comprises a state-of-the-art challenge for the field^86^.

Furthermore, although we observed no significant survival differences between HFD-fed and control-fed *Trp53*^*R270H/+*^ mice (Figure 2A), it remains possible that differences in tumor growth kinetics or aggressiveness exist but were not detectable due to the endpoint-based nature of our experimental design. It remains possible that effects of HFD-induced obesity on tumorigenesis or survival could emerge in a subset of animals with extended aging beyond this timeframe. Moreover, our assessment relied primarily on endpoint analysis and histological evaluation rather than continuous, longitudinal monitoring of tumor growth dynamics.

Beyond p53, this study identifies a new model for examining long-term tumor and host dynamics (genetic, transcriptional, immune, epigenetic and metabolic) in contexts of combined cancer susceptibility and metabolic disease evolution. The uncoupling of dietary effects from an altered cancer trajectory affords unique interdisciplinary opportunities to understanding chronic cancer and metabolic disease co-evolution (for up to ∼1yr).

## METHODS

### Origin and maintenance of mice

This research complies with ethical regulations and protocols approved by Institutional Animal Care and Use Committee (Van Andel Institute, USA; protocols 19-0026, 22-09-036, 18-10-028, and 21-08-023). *B6*.*129S4-Trp53<tm3*.*1Tyj>/J* (*Tp53*^*+/R270H*^) animals were originally generated in the Jacks lab^46^ and purchased from Jackson Laboratories (stock #008182). Mice were backcrossed for over 10 generations and maintained in house by breeding with wild-type siblings and periodic background refreshment using wild-types from JAX. 392 F1 hybrids were generated by crossing 8-week-old *FVB*.*J* males with two 8-week-old *B6*.*Tp53*^*R270H/+*^ females, which were separated after plug checking the next morning. Mating animals were randomly selected. All animals were fed breeder chow (Lab diet, 5021 cat. #0006540) *ad libitum* upon weaning and then randomly assigned to breeder chow or high-fat diet (Research diets, D12492i) at 8 weeks of age. Mice were housed in individually ventilated cages (Tecniplast, Sealsafe Plus GM500 in DGM Racks) at a density of maximum five animals per cage. Each cage was enriched with Enviro-dri (The Andersons, Crink-l’Nest) and cardboard dome homes (Shepherd, Shepherd Shack Dome). Whenever possible, same-sex siblings and same-sex animals from different litters were combined (∼20 days of age). Animals were kept on a 12-hour light/dark cycle at an average ambient temperature of 23°C and 35% humidity.

Body composition data was collected from 359 animals, including 173 males (74 WT and 99 *Tp53*^*+/R270H*^) and 186 females (82 WT and 104 *Tp53*^*+/R270H*^). Chow diet (CD, Lab diet, 5021 cat. #0006540) or high-fat diet (HFD, Research diets, D12492i) was randomly assigned at 8 weeks of age, as follows: 24 WT males under CD, 29 WT males under HFD, 38 *Tp53*^*+/R270H*^ males under CD, 56 *Tp53*^*+/R270H*^ males under HFD, 30 WT females under CD, 40 WT females under HFD, 41 *Tp53*^*+/R270H*^ females under CD, and 60 *Tp53*^*+/R270H*^ females under HFD. At 4, 8, 16, 32, 40, 50, 60, and 70 weeks of age (or at euthanasia), mice were weighed and scanned via EchoMRI for fat and lean mass composition in the morning (EchoMRI™, EchoMRI™-100H).

Tumor analysis was conducted on 182 animals: 85 males (7 WT-CD, 10 WT-HFD, 26 *Tp53*^*+/R270H*^-CD, and 42 *Tp53*^*+/R270H*^-HFD) and 97 females (8 WT-CD, 13 WT-HFD, 30 *Tp53*^*+/R270H*^-CD, and 46 *Tp53*^*+/R270H*^-HFD). We performed tumor analysis blinded for genotype and phenotype, temporally collecting mice according to the timing of health reports. We specify in the text every time we are only referring to one of the sexes.

### Statistics and reproducibility

Power analysis was performed by the Van Andel Institute (VAI) Bioinformatics and Biostatistics Core using the pwr R package for Power Analysis (R v.3.5.2)^87^, to determine sample size. In particular a 2 sample test of proportions was run based on an estimated effect size calculated on published data^46^ using a firth logistic regression. Power was set to 80%, alpha = 0.05, and assuming each group having equal sample sizes, considering 2 treatments (CD and HFD) and 5 different types of cancer evaluated (carcinoma, sarcoma, lymphoma, and leukemia).

Due to COVID-related reductions, 165 animals were randomly excluded for tumor analysis. Additionally, 7 mice died after birth, precluding further analysis. 10 were found dead and too stiff to harvest. The final cohort included all animals from litters of 5-12 pups.

Experiments were randomized, and investigators were blinded to group allocation and outcome assessment wherever possible.

### Genotyping

Ear punch biopsies were collected at 10 days and digested in 20 µl genomic DNA lysis buffer (100 mM Tris-HCl pH 8.5, 5 mM EDTA, 0.2% SDS, 100 mM NaCl) with 20 mg proteinase K (Thermo Scientific, EO0491). The thermal cycling protocol used was 55°C for 16 hours, 95°C for 10’, and a 4°C hold (lid at 105°C). Nuclease-free water (Invitrogen, AM9938) was added to each lysate for a final volume of 180 µl. PCR reactions for *Tp53* allele used 1 µl diluted biopsy lysate in a 19 µl master mix (1X DreamTaq Buffer, 0.2 mM dNTPs, 0.1 µM primer forward and reverse mix, 2 U DreamTaq DNA Polymerase, in nuclease-free water; Thermo Scientific, EP0703). PCR primer and thermal cycling conditions are detailed in Supplementary Tables 1 and 2. 20 µl of each PCR product were digested with 0.5 µl MslI (for *Tp53*^*R270H*/+^; New England BioLabs, R0571L) in a final reaction volume of 30 µl. Restriction conditions are detailed in Supplementary Table 3. Digestion products (∼500 bp WT *Tp53*, ∼200 + ∼300 bp *Tp53*^*R270H/+*^) were visualized on a 3% agarose gel (Fisher Scientific, BP160-500) in 1X TAE, with GelRed as intercalating dye (Biotium, 41003).

### Health monitoring

VAI Vivarium Core staff monitored mice 2-3 times per week for health, well-being, and abnormal mass/tumor presence. Mice were euthanized if they exhibited >20% weight loss, tumors ∼15% of body weight (this maximal tumor size was never exceeded), tumor ulcerations, tumor discharge or hemorrhage, mobility issues, reduced appetite or hydration, limited defecation or urination, abnormal gait or posture, labored breathing, lack of movement, or hypothermia. Mice with reported health concerns or those reaching the 70-week study endpoint were euthanized via CO_2_ asphyxiation and cervical dislocation.

### Tissue harvesting

Tissues were dissected and fixed in 10% NBF solution (3.7-4% formaldehyde 37-40%, 0.03 M NaH_2_PO_4_, 0.05 M Na_2_HPO_4_, in distilled water with final pH of 7.2± 0.5): epidydimal white adipose tissue (eWAT); uterus or preputial glands, seminal vesicles, and testis; bladder; pancreas; spleen; intestine; stomach; mesenteric fat; liver; kidneys; heart; lungs; thymus; brain; breast (9^th^); hindlimb muscles and bones. We also recovered spine, ribs, skull, skin, and any other mass if abnormal. Fixative volume was 15-20 times the tissue volume. Specimens > 2.5 mm thick were cut to proper fixation. Most tissues were fixed for 40 hours, while fat-rich tissues (eWAT, mesenteric fat, uterus) were fixed for 72 hours. Bones and spines were fixed for 1 week followed by 1-week decalcification in 14% EDTA (14% free-acid EDTA at pH 7.2, adjusted with NH_4_OH). After incubation, all tissues were moved to 70% ethanol. Data collection was blinded.

### Tissue preparation for histology

All tissues were paraffin-embedded by the VAI Pathology and Biorepository Core. Dehydration and clearing were automated with a Tissue-Tek VIP 5 (Sakura) using the following protocol: 60’ in 70% ethanol; 60’ in 80% ethanol; 2x 60’ in 95% ethanol; 3x 60’ in 100% ethanol; 2x 30’ in xylene; and 75’in paraffin. Embedding was performed with a Leica EG1150. Three 5-µm sections, spaced 150 µm apart, were cut from each tissue for hematoxylin and eosin (H&E) staining using a Leica rotary microtome. The remaining tissue was stored as a paraffin block. H&E staining was performed with a Tissue-Tek Prisma Plus Automated Slide Stainer (Sakura) and Prisma H&E Staining Kit #1.

### Pathology evaluation

Standard 5-µm H&E-stained sections were assessed for tumors and dysplastic lesions by a board-certified pathologist at the VAI Pathology and Biorepository Core. Most samples were provided blindly. Tumors were classified as malignant or benign, with all malignant tumors being primary. Metastatic or secondary tumors were identified based on primary tumor characteristics and immunohistochemical validation but were not reported in this study. Tumors were categorized into carcinomas, germ cell tumors, leukemias, lymphomas, and sarcomas, with detailed classification by tissue of origin.

## ACKNOWLEDGEMENTS

We thank E. Lien and D. Schramek for suggestions and support; as well as the MPI-IE Facilities and VAI Vivarium (RRID:SCR_023211), Transgenics (RRID:SCR_022914), Pathology and Biorepository (RRID:SCR_022912), and Bioinformatics and Biostatistics (RRID:SCR_024762) Cores. This work was supported by funding from VAI through internal philanthropy to JAP, NIH award number 1R01HG012444 to JAP, MeNU pilot project grants and Human Frontier Science Program Long-Term Fellowship LT000441/2018-L to IP. The funders had no role in study design, data collection and analysis, decision to publish or preparation of the manuscript.

## AUTHOR CONTRIBUTIONS

IP and JAP conceived the project, designed the overall methodology, and supervised the work. IP designed each individual experiment, performed the *in vivo* experiments, genotyping and the tissue harvesting together with MT. IP analyzed all the phenotypic data, while LF and SA supported data analysis. GH performed all the pathology reviews. IP and ZM performed statistical analyses. IP and JAP wrote, reviewed and edited the original draft. IP prepared the figures. JAP and IP acquired funds. JAP provided resources for the experiments.

## COMPETING INTERESTS STATEMENT

The authors declare no competing interests.

## FIGURE CAPTIONS

**Supplementary Figure 1.**
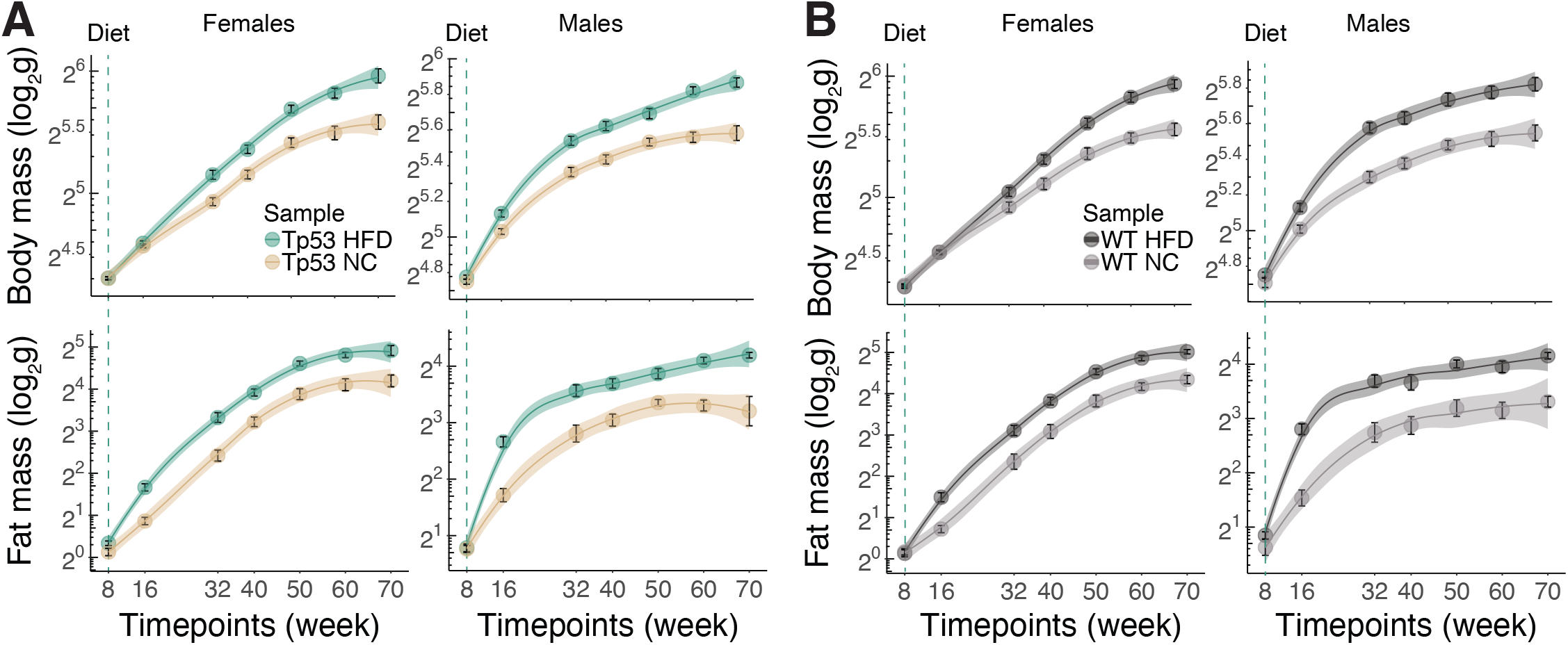
A cohort to study chronic effects of obesity on *Tp53*-dependent cancers across females and males. **A)** Scatter plots and smoothed conditional means (95% confidence interval, “loess” method) for *body* (<u>top</u>) and *fat* (<u>bottom</u>) mass in *Tp53*^*+/R270H*^ *females* (<u>left</u>) and *males* (<u>right</u>). N=101 females and 94 males (60 under high-fat and 41 under chow diet) and 94 males (56 under high-fat and 38 under chow diet). **B)** Scatter plots and smoothed conditional means (95% confidence interval, “loess” method) for *body* (<u>top</u>) and *fat* (<u>bottom</u>) mass in wild-type *females* (<u>left</u>) and *males* (<u>right</u>). N=70 females (40 under high-fat and 30 under chow diet) and 53 males (29 under high-fat and 24 under chow diet).

**Supplementary Figure 2.**
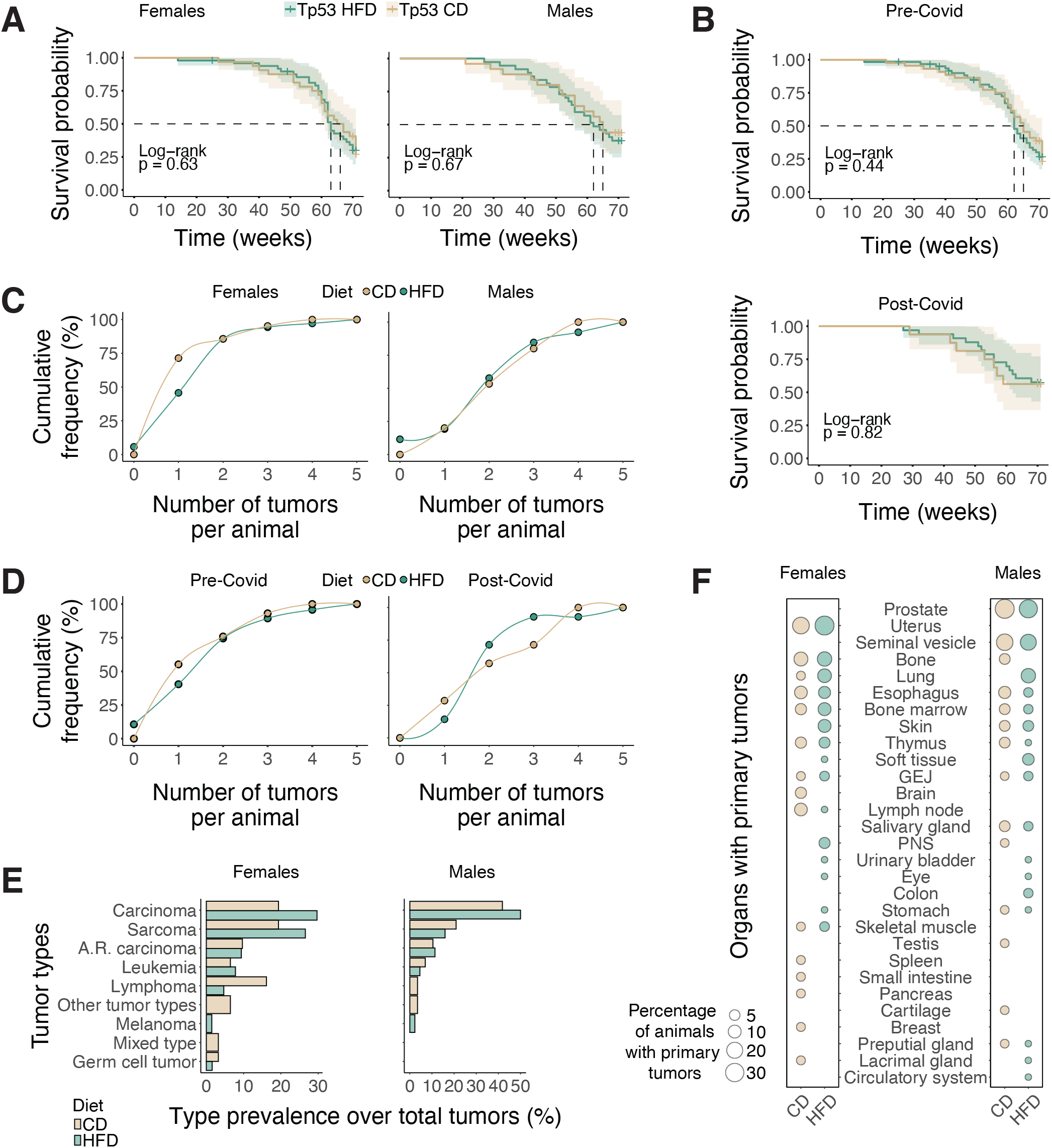
HFD does not alter survival, burden, or spectrum in *Tp53*^*R270H/+*^ female or male mice. **A)** Kaplan-Meier survival probability by diet for *Tp53*^*+/R270H*^ *females* (<u>left</u>) or *males* (<u>right</u>). Log-rank test, p=0.64 and p=0.67, respectively. N=81 females (49 high-fat diet vs 32 chow diet) and 74 males (46 high-fat diet vs 28 chow diet). **B)** Kaplan-Meier survival probability by diet for *Tp53*^*+/R270H*^ animals in the *pre-Covid* (<u>left</u>) or *post-Covid* (<u>right</u>) cohorts. Log-rank test, p=0.44 and p=0.82, respectively. N=106 pre-Covid (62 high-fat vs 44 chow diet) and 49 post-Covid (33 high-fat vs 16 chow diet). **C)** Cumulative distribution of tumor burden (number of tumors per animal) in *Tp53*^*+/R270H*^ *females* (<u>left</u>) or *males* (<u>right</u>) fed with chow– or high-fat diet. No significant differences were observed according to a two-sided, two-sample Kolmogorov-Smirnov test for either females (asymptotic; D = 0.037895, p-value = 1) or males (exact; D = 0.039722, p-value = 0.9469). N=56 females (35 high-fat diet vs 21 chow diet) and 41 males (26 high-fat vs 15 chow diet). **D)** Cumulative distribution of tumor burden (number of tumors per animal) in *Tp53*^*+/R270H*^ *pre-Covid* (<u>left</u>) or *post-Covid* (<u>right</u>) fed with chow– or high-fat diet. No significant differences were observed according to a two-sided, two-sample Kolmogorov-Smirnov test for either pre-Covid (asymptotic; D = 0.025907, p-value = 1) or post-Covid (exact; D = 0.041538, p-value = 0.9889). N=106 pre-Covid (62 high-fat vs 44 chow diet) and 49 post-Covid (33 high-fat vs 16 chow diet). **E)** Prevalence of each tumor type over the total number of tumors in chow– or high-fat diet-fed *Tp53*^*+/R270H*^ animals. N=95 tumors in females (64 in high-fat diet vs 31 in chow diet) and 73 in males (44 in high-fat diet vs 29 in chow diet). **F)** Percentage of *Tp53*^*+/R270H*^ animals with primary tumors targeting the different organs. N=54 females (33 high-fat diet vs 21 chow diet) and 38 males (23 high-fat diet vs 15 chow diet).

## Notes

### Competing Interest Statement

The authors have declared no competing interest.

### Summary of Updates

In this version, the manuscript has been updated to reflect important clarifications and refinements in terminology, data reporting, and interpretation. We provide more context about publication bias in scientific literature, particularly regarding negative results, and its impact on research priorities and patient care. The methods section now includes data validated in two independent cohorts conducted approximately two years apart, totaling 359 mice, and detailed statistical analyses. We also substantially expanded the discussion to explore potential mechanisms by which the *Tp53^R270H/+^ * mutation might counteract adverse metabolic and inflammatory changes typically associated with obesity. We've also added references supporting the role of p53 in regulating adipose tissue inflammation, immune cell infiltration, and adipokine signaling. Overall, these revisions enhance the clarity and rigor of the manuscript.

## REFERENCES

1 Ward, Z. J., Bleich, S. N., Long, M. W. & Gortmaker, S. L. Association of body mass index with health care expenditures in the United States by age and sex. PLOS ONE 16, e0247307 (2021). 10.1371/journal.pone.0247307

2 Renehan, A. G., Tyson, M., Egger, M., Heller, R. F. & Zwahlen, M. Body-mass index and incidence of cancer: a systematic review and meta-analysis of prospective observational studies. The Lancet 371, 569–578 (2008). 10.1016/S0140-6736(08)60269-X

3 Bhaskaran, K. et al. Body-mass index and risk of 22 specific cancers: a population-based cohort study of 5–24 million UK adults. The Lancet 384, 755–765 (2014). 10.1016/S0140-6736(14)60892-8

4 Moore, L. L., Chadid, S., Singer, M. R., Kreger, B. E. & Denis, G. V. Metabolic Health Reduces Risk of Obesity-Related Cancer in Framingham Study Adults. Cancer Epidemiology, Biomarkers & Prevention 23, 2057–2065 (2014). 10.1158/1055-9965.Epi-14-0240

5 Mahamat-saleh, Y. et al. Association of metabolic obesity phenotypes with risk of overall and site-specific cancers: a systematic review and meta-analysis of cohort studies. British Journal of Cancer (2024). 10.1038/s41416-024-02857-7

6 Lauby-Secretan, B. et al. Body Fatness and Cancer--Viewpoint of the IARC Working Group. N Engl J Med 375, 794–798 (2016). 10.1056/NEJMsr1606602

7 Marinac, C. R. et al. Body mass index throughout adulthood, physical activity, and risk of multiple myeloma: a prospective analysis in three large cohorts. Br J Cancer 118, 1013–1019 (2018). 10.1038/s41416-018-0010-4

8 Petrelli, F. et al. Association of Obesity With Survival Outcomes in Patients With Cancer: A Systematic Review and Meta-analysis. JAMA Network Open 4, e213520–e213520 (2021). 10.1001/jamanetworkopen.2021.3520

9 Ligibel, J. A. et al. American Society of Clinical Oncology position statement on obesity and cancer. J Clin Oncol 32, 3568–3574 (2014). 10.1200/jco.2014.58.4680

10 Rock, C. L. et al. Nutrition and physical activity guidelines for cancer survivors. CA: A Cancer Journal for Clinicians 62, 242–274 (2012). 10.3322/caac.21142

11 Senkus, E. et al. Primary breast cancer: ESMO Clinical Practice Guidelines for diagnosis, treatment and follow-up†. Annals of Oncology 24, vi7–vi23 (2013). 10.1093/annonc/mdt284

12 Font-Burgada, J., Sun, B. & Karin, M. Obesity and Cancer: The Oil that Feeds the Flame. Cell Metabolism 23, 48–62 (2016). 10.1016/j.cmet.2015.12.015

13 Perry, R. J. & Shulman, G. I. Mechanistic Links between Obesity, Insulin, and Cancer. Trends in Cancer 6, 75–78 (2020). 10.1016/j.trecan.2019.12.003

14 Li, R. et al. Transcriptome and DNA Methylome Analysis in a Mouse Model of Diet-Induced Obesity Predicts Increased Risk of Colorectal Cancer. Cell Reports 22, 624–637 (2018). 10.1016/j.celrep.2017.12.071

15 Calabrese, C. et al. Lipids and adipocytes involvement in tumor progression with a focus on obesity and diet. Obes Rev, e13833 (2024). 10.1111/obr.13833

16 Engin, A. B. & Engin, A. Next-Cell Hypothesis: Mechanism of Obesity-Associated Carcinogenesis. Adv Exp Med Biol 1460, 727–766 (2024). 10.1007/978-3-031-63657-8_25

17 Lysaght, J. & Conroy, M. J. The multifactorial effect of obesity on the effectiveness and outcomes of cancer therapies. Nature Reviews Endocrinology (2024). 10.1038/s41574-024-01032-5

18 Lennon, H., Sperrin, M., Badrick, E. & Renehan, A. G. The Obesity Paradox in Cancer: a Review. Current Oncology Reports 18, 56 (2016). 10.1007/s11912-016-0539-4

19 Nissen, S. B., Magidson, T., Gross, K. & Bergstrom, C. T. Publication bias and the canonization of false facts. eLife 5, e21451 (2016). 10.7554/eLife.21451

20 Fanelli, D. Negative results are disappearing from most disciplines and countries. Scientometrics 90, 891–904 (2011).

21 Dwan, K., Gamble, C., Williamson, P. R. & Kirkham, J. J. Systematic Review of the Empirical Evidence of Study Publication Bias and Outcome Reporting Bias — An Updated Review. PLOS ONE 8, e66844 (2013). 10.1371/journal.pone.0066844

22 Herrmann, D. et al. Statistical controversies in clinical research: publication bias evaluations are not routinely conducted in clinical oncology systematic reviews. Annals of Oncology 28, 931–937 (2017). 10.1093/annonc/mdw691

23 Easterbrook, P. J., Gopalan, R., Berlin, J. A. & Matthews, D. R. Publication bias in clinical research. The Lancet 337, 867–872 (1991). 10.1016/0140-6736(91)90201-Y

24 Mlinarić, A., Horvat, M. & Šupak Smolčić, V. Dealing with the positive publication bias: Why you should really publish your negative results. Biochem Med (Zagreb) 27, 030201 (2017). 10.11613/bm.2017.030201

25 Metcalfe, S. et al. Trastuzumab: possible publication bias. The Lancet 371, 1646–1648 (2008). 10.1016/S0140-6736(08)60706-0

26 Dickersin, K. & Min, Y. I. NIH clinical trials and publication bias. Online J Curr Clin Trials Doc No 50, [4967 words; 4953 paragraphs] (1993).

27 Brazil, R. Illuminating ‘the ugly side of science’: fresh incentives for reporting negative results. Nature (2024).

28 Bradley, S. H. et al. Reducing bias and improving transparency in medical research: a critical overview of the problems, progress and suggested next steps. J R Soc Med 113, 433–443 (2020). 10.1177/0141076820956799

29 Echevarría, L., Malerba, A. & Arechavala-Gomeza, V. Researcher’s Perceptions on Publishing “Negative” Results and Open Access. Nucleic Acid Therapeutics 31, 185–189 (2020). 10.1089/nat.2020.0865

30 Dickersin, K.Min, Y.-I. & Meinert, C. L. Factors Influencing Publication of Research Results: Follow-up of Applications Submitted to Two Institutional Review Boards. JAMA 267, 374–378 (1992). 10.1001/jama.1992.03480030052036

31 Thornton, A. & Lee, P. Publication bias in meta-analysis: its causes and consequences. Journal of Clinical Epidemiology 53, 207–216 (2000). 10.1016/S0895-4356(99)00161-4

32 Hines, R. B. et al. Effect of comorbidity and body mass index on the survival of African-American and Caucasian patients with colon cancer. Cancer 115, 5798–5806 (2009). 10.1002/cncr.24598

33 Navarro, W. H. et al. Effect of Body Mass Index on Mortality of Patients with Lymphoma Undergoing Autologous Hematopoietic Cell Transplantation. Biology of Blood and Marrow Transplantation 12, 541–551 (2006). 10.1016/j.bbmt.2005.12.033

34 Parker, A. S. et al. Greater body mass index is associated with better pathologic features and improved outcome among patients treated surgically for clear cell renal cell carcinoma. Urology 68, 741–746 (2006). 10.1016/j.urology.2006.05.024

35 Schlesinger, S. et al. Postdiagnosis body mass index and risk of mortality in colorectal cancer survivors: a prospective study and meta-analysis. Cancer Causes & Control 25, 1407–1418 (2014). 10.1007/s10552-014-0435-x

36 Hakimi, A. A. et al. An Epidemiologic and Genomic Investigation Into the Obesity Paradox in Renal Cell Carcinoma. JNCI: Journal of the National Cancer Institute 105, 1862–1870 (2013). 10.1093/jnci/djt310

37 Amptoulach, S., Gross, G. & Kalaitzakis, E. Differential impact of obesity and diabetes mellitus on survival after liver resection for colorectal cancer metastases. Journal of Surgical Research 199, 378–385 (2015). 10.1016/j.jss.2015.05.059

38 Brunner, A. M. et al. Association between baseline body mass index and overall survival among patients over age 60 with acute myeloid leukemia. American Journal of Hematology 88, 642–646 (2013). 10.1002/ajh.23462

39 Banack, H. R. & Kaufman, J. S. The obesity paradox: Understanding the effect of obesity on mortality among individuals with cardiovascular disease. Preventive Medicine 62, 96–102 (2014). 10.1016/j.ypmed.2014.02.003

40 Mayeda, E. R. & Glymour, M. M. The Obesity Paradox in Survival after Cancer Diagnosis: Tools for Evaluation of Potential Bias. Cancer Epidemiology, Biomarkers & Prevention 26, 17–20 (2017). 10.1158/1055-9965.Epi-16-0559

41 Park, Y., Peterson, L. L. & Colditz, G. A. The Plausibility of Obesity Paradox in Cancer—Point. Cancer Research 78, 1898–1903 (2018). 10.1158/0008-5472.Can-17-3043

42 Petrelli, F. et al. Obesity paradox in patients with cancer: A systematic review and meta-analysis of 6,320,365 patients. medRxiv, 2020.2004.2028.20082800 (2020). 10.1101/2020.04.28.20082800

43 Hollstein, M., Sidransky, D., Vogelstein, B. & Harris, C. C. p53 Mutations in Human Cancers. Science 253, 49–53 (1991). doi:10.1126/science.1905840

44 Jr., F. P. L. a. J. F. F. Soft-Tissue Sarcomas, Breast Cancer, and Other Neoplasms. Annals of Internal Medicine 71, 747–752 (1969). 10.7326/0003-4819-71-4-747 %m 5360287

45 Guha, T. & Malkin, D. Inherited TP53 Mutations and the Li–Fraumeni Syndrome. Cold Spring Harbor Perspectives in Medicine 7 (2017). 10.1101/cshperspect.a026187

46 Olive, K. P. et al. Mutant p53 gain of function in two mouse models of Li-Fraumeni syndrome. Cell 119, 847–860 (2004). 10.1016/j.cell.2004.11.004

47 Wang, C.-Y. & Liao, J. K. in mTOR: Methods and Protocols (ed Thomas Weichhart) 421–433 (Humana Press, 2012).

48 Buettner, R., Schölmerich, J. & Bollheimer, L. C. High-fat Diets: Modeling the Metabolic Disorders of Human Obesity in Rodents. Obesity 15, 798–808 (2007). 10.1038/oby.2007.608

49 George, V., Tremblay, A., Després, J. P., Leblanc, C. & Bouchard, C. Effect of dietary fat content on total and regional adiposity in men and women. Int J Obes 14, 1085–1094 (1990).

50 Saris, W. H. M. et al. Randomized controlled trial of changes in dietary carbohydrate/fat ratio and simple vs complex carbohydrates on body weight and blood lipids: the CARMEN study. International Journal of Obesity 24, 1310–1318 (2000). 10.1038/sj.ijo.0801451

51 Tucker, L. A. & Kano, M. J. Dietary fat and body fat: a multivariate study of 205 adult females. The American Journal of Clinical Nutrition 56, 616–622 (1992). 10.1093/ajcn/56.4.616

52 Rohm, T. V., Meier, D. T., Olefsky, J. M. & Donath, M. Y. Inflammation in obesity, diabetes, and related disorders. Immunity 55, 31–55 (2022). 10.1016/j.immuni.2021.12.013

53 Garaulet, M. & Madrid, J. A. Chronobiological aspects of nutrition, metabolic syndrome and obesity. Adv Drug Deliv Rev 62, 967–978 (2010). 10.1016/j.addr.2010.05.005

54 Adams, K. F. et al. Overweight, obesity, and mortality in a large prospective cohort of persons 50 to 71 years old. N Engl J Med 355, 763–778 (2006). 10.1056/NEJMoa055643

55 Dalgaard, K. et al. Trim28 Haploinsufficiency Triggers Bi-stable Epigenetic Obesity. Cell 164, 353–364 (2016). 10.1016/j.cell.2015.12.025

56 Yang, C.-H. et al. Independent phenotypic plasticity axes define distinct obesity sub-types. Nature Metabolism 4, 1150–1165 (2022). 10.1038/s42255-022-00629-2

57 Beyaz, S. et al. High-fat diet enhances stemness and tumorigenicity of intestinal progenitors. Nature 531, 53–58 (2016). 10.1038/nature17173

58 Park, E. J. et al. Dietary and Genetic Obesity Promote Liver Inflammation and Tumorigenesis by Enhancing IL-6 and TNF Expression. Cell 140, 197–208 (2010). 10.1016/j.cell.2009.12.052

59 Philip, B. et al. A high-fat diet activates oncogenic Kras and COX2 to induce development of pancreatic ductal adenocarcinoma in mice. Gastroenterology 145, 1449–1458 (2013). 10.1053/j.gastro.2013.08.018

60 Maguire, O. A. et al. Creatine-mediated crosstalk between adipocytes and cancer cells regulates obesity-driven breast cancer. Cell Metabolism 33, 499–512.e496 (2021). 10.1016/j.cmet.2021.01.018

61 Labbé, D. P. et al. High-fat diet fuels prostate cancer progression by rewiring the metabolome and amplifying the MYC program. Nature Communications 10, 4358 (2019). 10.1038/s41467-019-12298-z

62 Xia, S. et al. Prevention of Dietary-Fat-Fueled Ketogenesis Attenuates BRAF V600E Tumor Growth. Cell Metabolism 25, 358–373 (2017). 10.1016/j.cmet.2016.12.010

63 Panzeri, I. et al. TRIM28-dependent developmental heterogeneity determines cancer susceptibility through distinct epigenetic states. Nature Cancer 6, 385–403 (2025). 10.1038/s43018-024-00900-3

64 Berkers, Celia R., Maddocks Oliver D. K., Cheung Eric C., Mor, I. & Vousden, Karen H. Metabolic Regulation by p53 Family Members. Cell Metabolism 18, 617–633 (2013). 10.1016/j.cmet.2013.06.019

65 Khaddour, K., Gomez-Perez, S. L., Jain, N., Patel, J. D. & Boumber, Y. Obesity, Sarcopenia, and Outcomes in Non-Small Cell Lung Cancer Patients Treated With Immune Checkpoint Inhibitors and Tyrosine Kinase Inhibitors. Front Oncol 10, 576314 (2020). 10.3389/fonc.2020.576314

66 Zhang, X., Liu, Y., Shao, H. & Zheng, X. Obesity Paradox in Lung Cancer Prognosis: Evolving Biological Insights and Clinical Implications. J Thorac Oncol 12, 1478–1488 (2017). 10.1016/j.jtho.2017.07.022

67 Assumpção, J. A. F., Pasquarelli-do-Nascimento, G., Duarte, M. S. V., Bonamino, M. H. & Magalhães, K. G. The ambiguous role of obesity in oncology by promoting cancer but boosting antitumor immunotherapy. Journal of Biomedical Science 29, 12 (2022). 10.1186/s12929-022-00796-0

68 Tanti, J.-F., Ceppo, F., Jager, J. & Berthou, F. Implication of inflammatory signaling pathways in obesity-induced insulin resistance. Frontiers in Endocrinology 3 (2013). 10.3389/fendo.2012.00181

69 Heilbronn, L. K. & Campbell, L. V. Adipose tissue macrophages, low grade inflammation and insulin resistance in human obesity. Curr Pharm Des 14, 1225–1230 (2008). 10.2174/138161208784246153

70 Doerner, S. K. et al. High-Fat Diet-Induced Complement Activation Mediates Intestinal Inflammation and Neoplasia, Independent of Obesity. Molecular Cancer Research 14, 953–965 (2016). 10.1158/1541-7786.Mcr-16-0153

71 Wang, H. et al. An AMPK-dependent, non-canonical p53 pathway plays a key role in adipocyte metabolic reprogramming. eLife 9, e63665 (2020). 10.7554/eLife.63665

72 Al-Massadi, O. et al. Pharmacological and Genetic Manipulation of p53 in Brown Fat at Adult But Not Embryonic Stages Regulates Thermogenesis and Body Weight in Male Mice. Endocrinology 157, 2735–2749 (2016). 10.1210/en.2016-1209

73 Minamino, T. et al. A crucial role for adipose tissue p53 in the regulation of insulin resistance. Nature Medicine 15, 1082–1087 (2009). 10.1038/nm.2014

74 Liu, Y., Su, Z., Tavana, O. & Gu, W. Understanding the complexity of p53 in a new era of tumor suppression. Cancer Cell 42, 946–967 (2024). 10.1016/j.ccell.2024.04.009

75 Liu, Y. & Gu, W. The complexity of p53-mediated metabolic regulation in tumor suppression. Seminars in Cancer Biology 85, 4–32 (2022). 10.1016/j.semcancer.2021.03.010

76 Wang, P.-Y. et al. Increased Oxidative Metabolism in the Li–Fraumeni Syndrome. New England Journal of Medicine 368, 1027–1032 (2013). doi:10.1056/NEJMoa1214091

77 Li, T. et al. Tumor Suppression in the Absence of p53-Mediated Cell-Cycle Arrest, Apoptosis, and Senescence. Cell 149, 1269–1283 (2012). 10.1016/j.cell.2012.04.026

78 Wang, P.-Y. et al. Reducing Fatty Acid Oxidation Improves Cancer-free Survival in a Mouse Model of Li-Fraumeni Syndrome. Cancer Prevention Research 14, 31–40 (2021). 10.1158/1940-6207.Capr-20-0368

79 Wang, P.-y. et al. Inhibiting mitochondrial respiration prevents cancer in a mouse model of Li-Fraumeni syndrome. The Journal of Clinical Investigation 127, 132–136 (2017). 10.1172/JCI88668

80 Coral, D. E. et al. A phenome-wide comparative analysis of genetic discordance between obesity and type 2 diabetes. Nature Metabolism 5, 237–247 (2023). 10.1038/s42255-022-00731-5

81 Khera, A. V. et al. Polygenic Prediction of Weight and Obesity Trajectories from Birth to Adulthood. Cell 177, 587–596.e589 (2019). 10.1016/j.cell.2019.03.028

82 Cirulli, E. T. et al. Profound Perturbation of the Metabolome in Obesity Is Associated with Health Risk. Cell Metabolism 29, 488–500.e482 (2019). 10.1016/j.cmet.2018.09.022

83 Petersen, M. C. et al. Cardiometabolic characteristics of people with metabolically healthy and unhealthy obesity. Cell Metabolism 36, 745–761.e745 (2024). 10.1016/j.cmet.2024.03.002

84 Tabara, Y., Shoji-Asahina, A., Ogawa, A. & Sato, Y. Metabolically healthy obesity and risks of cardiovascular disease and all-cause mortality, a matched cohort study: the Shizuoka study. International Journal of Obesity 48, 1164–1169 (2024). 10.1038/s41366-024-01541-3

85 Schulze, M. B. & Stefan, N. Metabolically healthy obesity: from epidemiology and mechanisms to clinical implications. Nature Reviews Endocrinology (2024). 10.1038/s41574-024-01008-5

86 Bhaskaran, K. et al. Body-mass index and risk of 22 specific cancers: a population-based cohort study of 524 million UK adults. Lancet 384, 755–765 (2014). 10.1016/s0140-6736(14)60892-8

87 Champely, S., Ekstrom, C., Dalgaard, P., Gill, J., Weibelzahl, S., Anandkumar, A., Ford, C., Volcic, R., & De Rosario, H. pwr: Basic functions for power analysis. Software, < https://cran.rproject.org/web/packages/pwr/> (2017).

