## Supplementary Tables 1-3 for "Chronic obesity does not alter cancer incidence in *Trp53*^*R270H/*+^ mice"

### Table 1 – Genotyping primers.

| Target | Forward primer | Reverse primer |
|---|---|---|
| *Tp53* | ATGCGACTCTCCAGCCTTGGTA | TTGGGCTTAGGGACGTCTCTTATC |

### Table 2 – Genotyping PCR conditions.

| Target | Initial denaturation | Denaturation (x40 cycles) | Annealing (x40 cycles) | Extension (x40 cycles) | Final extension | Hold |
|---|---|---|---|---|---|---|
| *Tp53* | 95 °C for 3 minutes | 95 °C for 30 seconds | 63 °C for 30 seconds | 72 °C for 1 minute | 72 °C for 5 minutes | 4 °C infinite |

### Table 3 – Genotyping restriction conditions.

| Target | Digestion | Enzyme inactivation | Hold |
|---|---|---|---|
| *Tp53* | 37 °C for 30 minutes (MslI) | 80°C for 20 minutes | 4 °C infinite |
